# Evidence for recombination in dengue virus genomes

**DOI:** 10.64898/2026.06.14.732057

**Authors:** Hugo de Paula Oliveira, Denis Jacob Machado, Paula Prieto-Oliveira, Kary Ocaña

## Abstract

Recombination is a key driver of RNA virus evolution, yet its extent and evolutionary implications in dengue virus (DENV) remain incompletely understood. We conducted a comprehensive, genome-wide recombination screen across 6,905 complete DENV genomes representing all four serotypes, 82 countries, and eight decades of sampling (1944–2023) retrieved from the Bacterial and Viral Bioinformatics Resource Center. Using seven complementary recombination detection methods implemented in RDP5, we identified 66 recombination events across 53 unique recombinant sequences, of which 29 are newly described. Events included intra-genotypic (*n* = 18), inter-genotypic (*n* = 32), and inter-serotypic (*n* = 16) exchanges spanning 14 genotypes and four continents, with no meaningful serotype-level enrichment (Cramér’s *V* = 0.054). Recombination was concentrated in non-structural genes, most frequently *NS3* (19 events), *NS5* (17), and *NS2* (12), while the capsid gene contained no recombination events, consistent with strong functional constraint. Single-nucleotide polymorphism analyses confirmed low divergence between recombinants and their inferred parents in both recombinant and non-recombinant regions. Phylogenomic analysis of 6,642 sequences revealed that recombinants cluster significantly closer to their major parents (*p* = 8.9 × 10^−6^) and that their removal does not significantly alter tree topology (*p* = 0.898), suggesting that the short length of recombinant regions limits phylogenetic conflict. We also introduce RECOSIM, an unsupervised machine-learning tool for recombination detection that achieved higher precision than RDP5 on both simulated (93.4% vs. 80.0%) and empirical (98.1% vs. 39.3%) datasets. Collectively, these results establish recombination as a widespread, pan-serotypic phenomenon in DENV with implications for genomic surveillance, vaccine evaluation, and evolutionary inference.

## 1. Introduction

Dengue virus (DENV; *Flaviviridae*: *Orthoflavivirus*, species *Orthoflavivirus denguei*) is the causative agent of dengue fever (Harapan et al., 2020; Postler et al., 2023). The virus is transmitted by *Aedes* mosquitoes, which are widely distributed in tropical and subtropical regions (Diamond and Pierson, 2015). Clinical manifestations of DENV infection range from mild febrile illness to severe hemorrhagic syndrome and severe shock syndrome (Huy et al., 2013). As the most prevalent arthropod-borne viral disease worldwide, dengue causes hundreds of millions of infections annually, disproportionately affecting developing countries (Roy and Bhattacharjee, 2021). Dengue imposes substantial public health and economic burdens at both national and global scales (Shepard et al., 2016).

DENV virions are spherical, lipid-enveloped particles of approximately 50 nm in diameter. The capsid encloses a positive-sense single-stranded RNA (+ssRNA) genome of *∼*11 kb (Alcaraz-Estrada et al., 2010). The genome consists of a single open reading frame (ORF) flanked by structured 5’ and 3’ untranslated regions (King et al., 2008). The ORF encodes a single polyprotein that is subsequently cleaved into three structural proteins: capsid (C), premembrane (prM), and envelope (E); and seven nonstructural (NS) proteins: NS1, NS2A, NS2B, NS3, NS4A, NS4B, and NS5 (Chambers et al., 1990; Alcaraz-Estrada et al., 2010). Dengue viruses are classified into four antigenically distinct serotypes, DENV-1 to DENV-4, each further subdivided into genotypes (Hill et al., 2024). Phylogenetic analyses consistently recover tree topologies in which each serotype forms a well-supported monophyletic clade (Phadungsombat et al., 2018; Shrivastava et al., 2018). DENV genomes exhibit substantial inter-serotypic divergence, approximately 25–40% at the amino acid level, whereas inter-genotypic divergence within serotypes is typically below 10% (Holmes and Twiddy, 2003; Tamura et al., 2022), reflecting a well-documented history of lineage divergence.

In contrast to strict clonal evolution, recombination is a molecular mechanism in RNA viruses whereby genetic material is exchanged between co-infecting viruses, generating novel genomes (Pérez-Losada et al., 2015). It has been associated with viral evolution (Cebriá-Mendoza et al., 2023), host range expansion (Shahhosseini et al., 2021), and the emergence of new viruses (Hahn et al., 1988). Recombination has also been linked to increased virulence (Kononova et al., 2021), altered tissue tropisms (Hou et al., 2020), evasion of host immunity (Ritchie et al., 2014), and resistance to antivirals or vaccines (Shi et al., 2010). Detection of recombination is therefore required to avoid biases in phylogenetic analyses (Schierup and Hein, 2000; Martin et al., 2015). Undetected recombinants can yield inaccurate tree topologies and branch lengths, compromising evolutionary interpretations (Schierup and Hein, 2000; Ferretti et al., 2013).

Many tools are available for detecting recombination events, each implementing specific recombination detection methods (RDMs). RDMs employ phylogenetic methods (Milne et al., 2009; Samson et al., 2022), site pattern analysis (Bruen et al., 2006), pairwise sequence comparisons (Siepel et al., 1995; Samson et al., 2022), or population genetics (Kuhner, 2006; Guo et al., 2014). These methods collectively aim to identify genomic regions that originated through recombination. Once detected, recombination events can be further explored by comparing nucleotide similarity and polymorphism patterns between the recombinant sequence and its inferred parents. The major parent contributes the larger genomic fraction; the minor parent contributes the smaller fraction. Such comparisons characterize the genomic segments involved and reveal molecular signatures of recombination.

Dengue viruses have traditionally been considered to evolve primarily through the clonal accumulation of mutations (Chen and Vasilakis, 2011). Studies of recombination in DENV have focused on a limited number of genes and genomic datasets, and no empirical laboratory evidence has been documented (Holmes and Twiddy, 2003; Perez-Ramirez et al., 2009; Chen and Vasilakis, 2011; Sang et al., 2019). Nonetheless, recombination has been documented in other flaviviruses (Becher et al., 2001; Kalinina et al., 2002; Chuang and Chen, 2009), and DENV co-infections may support similar events (Dhanoa et al., 2016). Addressing this knowledge gap is relevant to public health, as recombination may contribute to the emergence of novel DENV variants and influence the design and safety evaluation of live-attenuated vaccines (Seligman and Gould, 2004; Monath et al., 2005; He et al., 2022; Singh et al., 2023).

In this study, we conducted a comprehensive genome-wide screening for recombination in DENV, examining its prevalence, genomic distribution, and evolutionary implications. We integrated large-scale phylogenomic analyses with detailed characterization of recombinant segments to elucidate the recombination landscape in DENV and to provide a framework for future studies of viral evolution and recombination detection.

## 2. Methods

### 2.1. Genomic data collection

A total of 6,905 complete human-isolated DENV genome sequences were retrieved from the Bacterial and Viral Bioinformatics Resource Center (https://www.bv-brc.org/). These genomes represent viruses sampled between 1944 and 2023 across 82 countries and were predominantly from Asia (4,928), followed by the Americas (1,798), Africa (57), Oceania (97), and Europe (9). The sequence information, including accession IDs, collection year, and country of isolation, is summarized in Table S1. Four complete genome sequences were sourced from the National Center for Biotechnology Information (https://www.ncbi.nlm.nih.gov/) as outgroups for the phylogenetic analysis: one Zika virus (MH882548) and three West Nile viruses (HM147823, GQ903680, and JX041632). Outgroup sequences were selected due to their close phylogenetic relationship to DENV.

### 2.2. Serotype and genotype classification

The Dengue Virus Typing Tool v4.2 (Vilsker et al., 2019) was used to determine the genotype and serotype of all 6,905 DENV sequences, using default parameters (Table S2; accessed on August, 2025). To accommodate computational tool constraints, the dataset was partitioned and processed in four independent, parallel batches

After serotype and genotype assignment, a chi-square test was conducted to compare the distribution of recombination events across serotypes, and Cramér’s V was calculated to assess the strength of association. Sequences had been previously classified as recombinant or non-recombinant using RDP5 and were grouped by serotype for analysis. The null hypothesis assumed equal distribution of recombination across serotypes.

### 2.3. Genome annotation and alignment

The 6,905 DENV complete genome sequences were submitted to the FLAVi (De Bernardi Schneider et al., 2020) pipeline for accurate gene annotation. FLAVi considers individual gene annotations to produce the final alignment, minimizing misalignments between genes. The pipeline was executed with default parameters (Table S2).

FLAVi returned two main outputs: (i) annotated and aligned sequences in FASTA format comprising 15,213 nt and (ii) a gene partition file in NEXUS format indicating the start and end positions of each gene within the alignment (Supplementary Data 3), allowing for the subsequent identification of recombination at the gene level.

### 2.4. Recombination detection

Recombination detection was performed on the 6,905 annotated and aligned DENV sequences using RDP v5.55 (Martin et al., 2021). Analyses were conducted with the options “Require topological evidence”, “Polish break-points”, “Check alignment consistency”, and “Genomes sequences as linear” (Table S2).

The dataset was initially screened for recombination signals using the RDP, GENECONV, and MaxChi methods, which are recommended in the RDP5 manual for the preliminary analysis of large datasets (*p*-value *<* 0.05). Candidate recombination signals were subsequently re-evaluated using the methods RDP, GENECONV, BootScan, MaxChi, Chimaera, SiScan, and 3Seq. Recombination events supported by three or more methods with a Bonferroni-corrected significance threshold of 0.05 were retained.

Detected recombination events were exported in CSV format, including estimated breakpoint positions, inferred major and minor parent sequences when present, and associated *p*-values. In total, 66 recombination events were identified, of which 51 had both parent sequences identified. The recombinant sequences presenting both minor and major parents comprised a dataset of 153 sequences, which was subsequently used for SNP analysis.

### 2.5. Recombinant SNPs analysis

A single-nucleotide polymorphism (SNP) analysis was conducted using the SNIPIT pipeline (O’Toole et al., 2024). The pipeline was executed using the command line described in Table S2, with the recombinant sequence defined as the reference. SNIPIT was applied to 51 FASTA datasets, each containing a recombinant genome and its inferred minor and major parents. The analysis generated a CSV file and graphical output identifying sequence variations relative to the reference. SNPs were examined in recombinant and non-recombinant regions.

Subsequently, sequence similarity between the recombinant genomes and their parents was assessed by visual inspection of RDP5 similarity plots, which compare sequences using 30 nt sliding windows. Seven recombination events were selected as case studies, representing different serotypes, genotypes, and recombinant region lengths and locations.

### 2.6. Literature search

A literature search was conducted in August 2025 to determine which recombination events are novel and which have been previously reported. Literature screening was performed in Google Scholar and PubMed, including peer-reviewed scientific articles and doctoral dissertations, with no publication date restriction. The accession IDs of the recombinant DENV sequences detected in this work were individually queried in both databases using: (dengue virus OR DENV) AND (recombination OR recombinant) AND ID.

Retrieved records were manually inspected to determine whether each queried sequence had been previously reported as recombinant. The inspection included the main text, supplementary materials, tables, and figures to verify recombinant classification and identify the assigned minor and major parents.

### 2.7. Phylogenetic analyses

#### Dataset

The initial dataset for phylogenetic analyses comprised 6,905 DENV, one ZIKV, and three WNV complete genome sequences, all annotated and aligned with FLAVi using default parameters. Identical sequences were identified and removed to reduce redundancy, while retaining all recombinants and their parents, resulting in 6,642 unique flavivirus sequences (Supplementary Data 4).

#### Tree construction

The phylogenomic tree topology was inferred under maximum likelihood using RAxML v8.2.12. Branch lengths were estimated under parsimony with TNT v1.6, and branch support was calculated using the Shimodaira-Hasegawa approximate likelihood ratio test (SH-aLRT) implemented in IQ-Tree v2.0.7. The tree inference pipeline is detailed in Figure S1.

#### Phylogenomic analyses

The effects of recombinant sequences on the phylogenomic tree were assessed. Differences in branch length distributions between recombinant and non-recombinant terminals were tested using the Mann–Whitney U test, and correlations between clade level recombination frequency and branch length were assessed using Spearman’s rank correlation. Recombinant proportions and branch lengths for phylogenetic clades are provided in Table S3. The phylogenetic proximity between recombinants and their inferred parents was evaluated using a custom script and the recombinant-parent assignments provided in Table S4. Statistical procedures are provided in Supplementary Data 4.

The contribution of each sequence to phylogenomic tree topology was evaluated using YBYRÁ (Jacob Machado, 2015), as described in Supplementary Data 4. Due to computational constraints, analyses were conducted on a pruned dataset retaining all recombinant sequences, their inferred parents, and up to three non-recombinant sequences per serotype and genotype, totaling 192 terminals. The pruned DENV phylogenomic tree was used as the reference and compared with alternative trees inferred after removal of single terminals from the alignment, allowing quantification of individual topological effects. Topological distances between each alternative tree and the reference tree were then calculated.

### 2.8. Execution and evaluation of RECOSIM

RECOSIM is an unsupervised machine-learning approach for recombination detection based on phylogenetic information. It requires prior categorization of character state transformations using YBYRÁ, representing nucleotide changes mapped onto tree branches. The resulting TSV file and the corresponding multiple sequence alignment in NEXUS format are used as input. Recombination signals are detected as clusters of categorized transformations. For each taxon, transformations are grouped using the DBSCAN algorithm, and overlapping clusters are filtered. Clusters from different taxa are then compared and overlapping regions that meet a predefined threshold are selected as candidates. Genomic regions corresponding to these clusters are extracted from the alignment and evaluated based on pairwise sequence identity. Regions exceeding a defined identity threshold are classified as recombinant. Implementation is available at https://gitlab.com/phyloinformatics/recosim.git, with additional details provided in Supplementary Data 5.

To evaluate the performance of RECOSIM, distinct tree topologies were used to represent tree-like evolutionary scenarios. Fifty empirical phylogenetic trees were obtained from studies deposited in the TreeBASE repository, while four simulated trees obtained from the random tree models available in IQ-TREE 2. These tree topologies were used to simulate sequence alignments with AliSim (implemented in IQ-TREE 2) under different evolutionary models (Table S5). Recombination was introduced by modifying sequences within each simulated alignment. For each event, a recombinant sequence and its corresponding minor and major parents were randomly assigned. A central genomic fragment from the minor parent replaced the corresponding region in the recombinant sequence. Similarly, flanking regions from the major parent replaced the corresponding regions in the recombinant.

RECOSIM was employed using three clustering strategies (regularClusters, expandingClusters, and expandingNotDeleting), which differ in how transformation clusters are merged and filtered (Supplementary Data 5). Multiple parameter combinations were tested, including epsilon (e), min_samples (m), overlap_threshold (o), and identity_threshold (i), which control clustering, overlap filtering, and sequence identity thresholds for recombination detection. Recombination detection perfor-mance was evaluated using hit and fail ratios, sensitivity, and precision (Supplementary Data 5), and compared with results obtained using RDP5 on the same datasets. Detection accuracy was defined as the proportion of correctly identified recombination events. RECOSIM was executed with regularClusters.py using the following parameters: o = 30, i = 0.95, e = 30, and m = 3.

## 3. Results and discussion

### 3.1. Serotyping and genotyping analysis of DENV genomes

The genomic dataset encompassed all four DENV serotypes: DENV-1 (2,910 sequences), DENV-2 (1,840), DENV-3 (1,436), and DENV-4 (719) (Table 1). Genotypic diversity spanned DENV-1 (genotypes I, II, III, IV, V, and VI), DENV-2 (I, II, III, V, VI), DENV-3 (I, II, III, V), and DENV-4 (I, II, III, and IV). A minor fraction of sequences (n = 27) remained unassigned to any specific genotype due to ambiguous phylogenetic support.

**Table 1.** Distribution of DENV genomes per serotype (Sero.) and genotype. Sequences with unassigned genotype by Genome Detective are labeled as ‘?’. T indicate line totals.

| Sero. | Genotype (#) |  |  |  |  |  |  | Total |
| --- | --- | --- | --- | --- | --- | --- | --- | --- |
|  | I | II | III | IV | V | VI | ? |  |
| DENV-1 | 2,186 | 6 | 212 | 83 | 416 | 4 | 3 | 2,910 |
| DENV-2 | 18 | 754 | 714 | – | 328 | 4 | 22 | 1,840 |
| DENV-3 | 95 | 622 | 714 | – | 4 | – | 1 | 1,436 |
| DENV-4 | 494 | 214 | 9 | 1 | – | – | 1 | 719 |

### 3.2. Identification of recombination events in DENV genomes with RDP5

Screening of the 6,905 aligned genomes yielded a total of 66 recombination events distributed across 53 unique recombinant sequences. Table 2 provides the ID, serotype, genotype, collection year, country of isolation, major and minor parents ID, gene, and breakpoints of each recombinant event identified.

**Table 2.** Recombination events detected in DENV genomes. (rows 1–32 of 66). Sequences are listed as ‘ID (Serotype-Genotype, Year, Country)’. Unknown parents are indicated by ‘—’. Carets (^) mark events in which one of the parents may itself be a recombinant. Asterisks (*) denote inter-serotypic or inter-genotypic events. Breakpoints are given in alignment coordinates. Sequences reported as recombinants for the first time are shown in bold

| Recombinant | Minor parent | Major parent | Gene(s) | Position |
| --- | --- | --- | --- | --- |
| FJ898399 (DENV-1-I, 2006, Viet Nam) | GQ199828 (DENV-1-I, 2007, Viet Nam) | GQ199806 (DENV-1-I, 2007, Viet Nam) | NS1 | 4,389–5,050 |
| JF459993 (DENV-1-I, 2002, Myanmar) | JN544411 (DENV-1-I, 2011, Singapore) | KT827371 (DENV-1-I, 2014, China) |  | Uncertain–4,573 |
| MN018333 (DENV-1-I, 2016, China) | JN544407 (DENV-1-I, 2011, Singapore) | MN018332 (DENV-1-I, 2016, China) | NS3 | 8,420–9,544 |
| **ON123666** (DENV-1-I, 2018, India) | MZ285732 (DENV-3-IV, 2019, India) | MF033196 (DENV-1-I, 2011, Singapore) | NS3, NS4A, NS4B | 8,682–10,890 |
| **ON123656** (DENV-1-III, 2019, India) | MZ285732 (DENV-3-IV, 2019, India) | MZ229305 (DENV-1-III, 2016, India) | NS2 | 5,979–6,490 |
| **ON123656** (DENV-1-III, 2019, India) | MZ285732 (DENV-3-IV, 2019, India) | ON123657 (DENV-1-III, 2019, India) | E | 2,964–3,617 |
| KU509261 (DENV-1-IV, 2010, Indonesia) | KU509260 (DENV-1-I, 2008, Cambodia) | JN544411 (DENV-1-IV, 2011, Singapore) | NS2, NS3 | 6,099–7,668 |
| **MW582813** (DENV-1-IV, 2019, China) | MZ285732 (DENV-3-IV, 2019, India) | MN018333 (DENV-1-I, 2016, China) |  | 5,371–Uncertain |
| **MW582813** (DENV-1-IV, 2019, China) | MZ285732 (DENV-3-IV, 2019, India) | OM281584 (DENV-1-I, 2019, Cambodia) | NS4B, NS5 | 11,183–12,711 |
| **MW582813** (DENV-1-IV, 2019, China) | MZ285732 (DENV-3-IV, 2019, India) | — | NS5 | 13,276–14,294 |
| **MW582813** (DENV-1-IV, 2019, China) | MZ285732 (DENV-3-IV, 2019, India) | OM281586 (DENV-1-I, 2019, Thailand) |  | Uncertain–4,618 |
| **MW582814** (DENV-1-IV, 2019, China) | MN018332 (DENV-1-I, 2016, China) | MZ285732 (DENV-3-IV, 2019, India) | NS2, NS3 | 6,628–8,726 |
| **MW582814** (DENV-1-IV, 2019, China) | MZ285732 (DENV-3-IV, 2019, India) | OM281584 (DENV-1-I, 2019, Cambodia) | E, NS1 | 3,456–4,683 |
| **MW582814** (DENV-1-IV, 2019, China) | MZ285732 (DENV-3-IV, 2019, India) | GQ199840 (DENV-1-I, 2006, Viet Nam) | NS4B, NS5 | 11,078–12,775 |
| **OM281589** (DENV-1-IV, 2019, Philippines) | OM281590 (DENV-1-I, 2019, Cambodia) | MZ285732 (DENV-3-IV, 2019, India) | NS5 | 13,962–14,312 |
| **OM281589** (DENV-1-IV, 2019, Philippines) | MF033260 (DENV-1-I, 2016, Singapore) | GQ868602 (DENV-1-IV, 2004, Philippines) | NS3 | 8,915–9,586 |
| **JN903578** (DENV-1-V, 2007, India) | JN903579 (DENV-1-I, 2008, India) | DQ285559 (DENV-1-V, 2004, Reunion) | NS1, NS2 | 5,415–6,912 |
| **KF184975** (DENV-1-VI, 2013, Angola) | MF033196 (DENV-1-I, 2011, Singapore) | GU131835 (DENV-1-V, 2004, Venezuela) |  | Uncertain–3,282 |
| GQ398257 (DENV-2-I, 1977, Indonesia) | GQ398269 (DENV-2-I, 1994, Puerto Rico) | GQ868588 (DENV-2-I, 1983, Mexico) | NS2, NS3, NS4A | 6,796–9,913 |
| JQ922552 (DENV-2-I, 1960, India) | JQ922548 (DENV-1-I, 2005, India) | KJ918750 (DENV-2-V, 2007, India) | NS5 | 12,148–12,575 |
| JQ922552 (DENV-2-I, 1960, India) | JQ922549 (DENV-2-I, 1996, India) | KJ918750 (DENV-2-V, 2007, India) | NS3, NS4A | 9,192–10,134 |
| JQ922553 (DENV-2-I, 1980, India) | KF041236 (DENV-2-I, 2008, Pakistan) | GQ398257 (DENV-2-I, 1977, Indonesia) | NS5 | 12,607–13,118 |
| **KC964094** (DENV-2-II, 1993, China) | KC964095 (DENV-2-V, 1998, China) | MT921572 (DENV-2-II, 2000, Australia) |  | Uncertain–1,836 |
| KU517845 (DENV-2-II, 2013, New Guinea) | KU517846 (DENV-2-I, 2014, Indonesia) | MT921570 (DENV-2-II, 2013, Australia) | E | 2,730–3,840 |
| **KX380821** (DENV-2-II, 2012, Singapore) | MN018357 (DENV-2-I, 2014, China) | KM279544 (DENV-2-II, 2012, Singapore) |  | Uncertain–3,348 |
| **KX380836** (DENV-2-II, 2013, Singapore) | KM279544 (DENV-2-I, 2012, Singapore) | MN018357 (DENV-2-II, 2014, China) | NS3, NS4A, NS4B | 8,746–11,424 |
| **KX452048** (DENV-2-II, 2014, Malaysia) | — | MK564483 (DENV-2-II, 2017, China) | E | 2,784–2,938 |
| KY937188 (DENV-2-II, 2015, China) | — | KY937190 (DENV-2-II, 2015, China) | pr, M | 1,165–1,540 |
| KY937189 (DENV-2-II, 2015, China) | — | KY937190 (DENV-2-II, 2015, China) |  | Uncertain–1,540 |
| **MF156233** (DENV-2-II, 2015, China) | KY586692 (DENV-2-V, 2001, Thailand) | MK578531 (DENV-2-II, 2016, China) | NS5 | 13,692–14,658 |
| **MF156235** (DENV-2-II, 2015, China) | KY586692 (DENV-2-V, 2001, Thailand) | MK578531 (DENV-2-II, 2016, China) | NS5 | 13,692–14,658 |
| **MF156244** (DENV-2-II, 2015, China) | GQ868591 (DENV-2-V, 1964, Thailand) | MK578531 (DENV-2-II, 2016, China) | NS5 | 13,696–14,026 |

Table 2. (continued)
| Recombinant | Minor parent | Major parent | Gene(s) | Position |
| --- | --- | --- | --- | --- |
| MG560143<sup>^</sup> (DENV-2-II, 2014, India) | MK783210 (DENV-2-I, 2018, China) | OM681318 (DENV-2-II, 2021, India) | E, NS1, NS2 | 2,426–6,691 |
| **MG560144** (DENV-2-II, 2014, India) | MG560143 (DENV-2-I, 2014, India) | MH822950 (DENV-2-II, 2013, India) | NS4B, NS5 | 10,874–12,197 |
| MH110575 (DENV-2-II, 2017, China) | MN018359 (DENV-2-I, 2017, China) | KY672953 (DENV-2-II, 2015, China) | NS3 | 8,675–9,218 |
| **ON123651** (DENV-2-II, 2020, India) | GQ868591 (DENV-2-V, 1964, Thailand) | MG721057 (DENV-2-II, 2016, India) | NS3, NS4A, NS4B | 9,318–11,100 |
| **EU482721** (DENV-2-III, 2006, USA) | FJ906956 (DENV-2-I, 2005, Nicaragua) | EU687213 (DENV-2-III, 2004, USA) | NS4B, NS5 | 11,098–13,898 |
| **EU482726** (DENV-2-III, 2006, USA) | KF955359 (DENV-2-I, 2006, Puerto Rico) | JX669477 (DENV-2-III, 2010, Brazil) | NS3, NS4A, NS4B, NS5 | 8,287–13,507 |
| GQ398269 (DENV-2-III, 1994, Puerto Rico) | GQ868591 (DENV-2-V, 1964, Thailand) | EU482562 (DENV-2-III, 1989, USA) | NS2, NS3 | 6,884–9,232 |
| GQ398269 (DENV-2-III, 1994, Puerto Rico) | GQ398301 (DENV-2-I, 1995, Puerto Rico) | GQ398281 (DENV-2-III, 1994, Puerto Rico) |  | Uncertain–13,232 |
| **GQ398282** (DENV-2-III, 1994, Puerto Rico) | GQ398301 (DENV-2-I, 1995, Puerto Rico) | GQ398283 (DENV-2-III, 1994, Puerto Rico) | NS3, NS4A, NS4B, NS5 | 9,204–13,194 |
| KF744399<sup>^</sup> (DENV-2-IV, 1998, Philippines) | KU509269 (DENV-2-I, 2009, Philippines) | KF744400 (DENV-2-IV, 2000, Philippines) | NS4A, NS4B | 10,083–11,210 |
| **KF744402<sup>^</sup>** (DENV-2-IV, 2000, Philippines) | MN566111 (DENV-2-I, 2018, New Caledonia) | KF744400 (DENV-2-IV, 2000, Philippines) | NS5 | 12,468–13,016 |
| **KF744402<sup>^</sup>** (DENV-2-IV, 2000, Philippines) | — | KF744399 (DENV-2-IV, 1998, Philippines) | E, NS1 | 3,750–4,606 |
| **EU482784** (DENV-2-V, 2003, Viet Nam) | MK783204 (DENV-2-I, 2018, China) | FM210216 (DENV-2-V, 2004, Viet Nam) | NS1, NS2 | 4,986–6,314 |
| **JN368476** (DENV-2-V, 2007, Cambodia) | KY586591 (DENV-2-V, 2002, Thailand) | FJ639717 (DENV-2-V, 2007, Cambodia) |  | 12,287–Uncertain |
| **KC964095** (DENV-2-V, 1998, China) | KC964094 (DENV-2-I, 1993, China) | KY586679 (DENV-2-V, 2001, Thailand) | NS5 | 12,938–14,092 |
| OP909734 (DENV-2-V, 2020, Singapore) | GQ868591 (DENV-2-V, 1964, Thailand) | — |  | 6,086–Uncertain |
| JF295012 (DENV-3-II, 2007, Cambodia) | ON103302 (DENV-1-I, 2021, Bangladesh) | FJ639725 (DENV-3-II, 2003, Cambodia) | NS3 | 8,430–8,820 |
| **KF955462<sup>^</sup>** (DENV-3-II, 2001, Cambodia) | — | MH891766 (DENV-3-III, 2017, India) |  | Uncertain–Uncertain |
| KY586804 (DENV-3-II, 2007, Thailand) | KY586785 (DENV-3-I, 1995, Thailand) | MW946661 (DENV-3-II, 2008, Thailand) | NS3, NS4A | 8,470–9,640 |
| **KY586818\*** (DENV-3-II, 2007, Thailand) | KY586814 (DENV-3-I, 2006, Thailand) | KY863456 (DENV-3-I, 2016, Indonesia) | E, NS1, NS2, NS3 | 3,152–8,446 |
| KY586820<sup>^</sup> (DENV-3-III, 2008, Thailand) | KY586721 (DENV-3-I, 2000, Thailand) | MW946767 (DENV-3-III, 2006, Thailand) |  | 14,684–Uncertain |
| **OP895705** (DENV-3-III, 2018, India) | FJ882602 (DENV-2-I, 1996, Sri Lanka) | OQ948474 (DENV-3-III, 2019, China) |  | 14,298–Uncertain |
| OQ948474 (DENV-3-III, 2019, China) | — | KX380841 (DENV-3-III, 2012, Singapore) |  | Uncertain–4,844 |
| **JN638572** (DENV-4-I, 2008, Cambodia) | OL314735 (DENV-4-I, 2019, Indonesia) | MW945520 (DENV-4-I, 2006, Thailand) | NS4B, NS5 | 11,267–12,451 |
| JQ922559<sup>^</sup> (DENV-4-I, 1979, India) | JF262783 (DENV-4-I, 1961, India) | MW945600 (DENV-4-I, 2005, Thailand) | E | 2,461–3,262 |
| JQ922559<sup>^</sup> (DENV-4-I, 1979, India) | MN018394 (DENV-4-I, 2015, China) | JQ922560 (DENV-4-I, 2009, India) | NS5 | 13,464–13,972 |
| MN018395 (DENV-4-I, 2015, China) | MN018396 (DENV-4-I, 2015, China) | MW945520 (DENV-4-I, 2006, Thailand) | NS3, NS4A | 8,354–10,007 |
| MN018395 (DENV-4-I, 2015, China) | MK640208 (DENV-4-II, 2018, China) | JN638572 (DENV-4-I, 2008, Cambodia) | NS1, NS2 | 5,302–6,620 |
| MN018395 (DENV-4-I, 2015, China) | MN018393 (DENV-4-I, 2015, China) | JN638572 (DENV-4-I, 2008, Cambodia) |  | Uncertain–1,188 |
| KU523871 (DENV-4-II, 2014, Philippines) | OL314735 (DENV-4-I, 2019, Indonesia) | KC333651 (DENV-4-II, 2012, China) | NS5 | 12,772–13,814 |
| **MH823210\*** (DENV-4-II, 2014, Indonesia) | MK783210 (DENV-2-I, 2018, China) | OM681318 (DENV-2-II, 2021, India) | E, NS1, NS2 | 2,426–6,691 |
| MK614089 (DENV-4-II, 2018, China) | MN018397 (DENV-4-I, 2016, China) | MK640208 (DENV-4-II, 2018, China) | NS2, NS3 | 6,612–7,652 |
| MN018396 (DENV-4-II, 2015, China) | MW945613 (DENV-4-I, 2014, Thailand) | MK640208 (DENV-4-II, 2018, China) | NS2, NS3 | 6,165–7,914 |
| **OL314740\*** (DENV-4-II, 2019, Indonesia) | MK783210 (DENV-2-I, 2018, China) | OM681318 (DENV-2-II, 2021, India) | E, NS1, NS2 | 2,426–6,691 |

Of the 66 recombination events, intra-genotypic events (18; same serotype and genotype), inter-genotypic events (32; same serotype, different genotypes), and interserotypic events (16; different serotypes and genotypes) were identified (Table 2). Among the recombinant sequences, GQ398269, JQ922552, JQ922559, KF744402, OM281589, and ON123656 harbored two distinct recombination events each, whereas genomes MN018395 and MW582814 exhibited three independent events each, and MW582813 presented four recombination events.

The occurrence of multiple recombination events within the same sequence was reported for other positive-sense RNA viruses, in which successive recombination contributes to increased viral diversification (Ngo et al., 2024). Interestingly, ON123656, MW582814, and MW582813 presented multiple recombination events involving the same minor parents, a pattern consistent with successive template-switching events during RNA replication (Chen et al., 2008). Alternatively, these multievent patterns may represent artificial fragmentations of a single, larger recombination event that the applied detection algorithms failed to resolve as a cohesive unit.

Inter-serotypic recombination has been regarded as rare in DENV (Domingo et al., 2006; Craig et al., 2003), and has been suggested to contribute to the emergence of disease-associated strains (Worobey et al., 1999). In contrast, 16 inter-serotypic events were identified in the present study. The detection of recombination events across distinct serotypes and diverse geographic regions underscores the importance of ongoing surveillance, as it could contribute to viral diversity and potentially impact virulence, transmission, or vaccine efficacy (Seligman and Gould, 2004).

A total of 53 unique recombinant DENV sequences were distributed across all four serotypes, with DENV-2 exhibiting the highest prevalence (27 sequences), followed by DENV-1 (11), DENV-4 (8), and DENV-3 (7) (Table 3). Recombinants were distributed among 14 genotypes, with DENV-2 genotype II being the most represented (14 sequences). Although DENV-2 accounted for 50.94% of the recombinant sequences and the chi-square test indicated a statistically significant non-uniform distribution among serotypes (*χ*^2^ = 20.24; *p* = 0.000152), the effect size was negligible (Cramér’s *V* = 0.054). Consequently, this disparity is an artifact of our large sample size rather than biological evidence of a preferential association between recombination mechanisms and any specific viral serotype.

**Table 3.** Distribution of recombinant DENV genomes per serotype (Sero.) and genotype. # indicates the number of recombinants; T indicates line totals.

| Sero. | Genotype (#) |  |  |  |  |  | Total |
| --- | --- | --- | --- | --- | --- | --- | --- |
|  | I | II | III | IV | V | VI |  |
| DENV-1 | 4 | — | 1 | 4 | 1 | 1 | 11 |
| DENV-2 | 3 | 14 | 4 | 2 | 4 | — | 27 |
| DENV-3 | — | 4 | 3 | — | — | — | 7 |
| DENV-4 | 3 | 5 | — | — | — | — | 8 |

Geographically, most recombinant sequences were detected in Asia (47 of 53), followed by North America (4), Africa (1), and Oceania (1) (Table S6). China (15) and India (10) were the countries with the highest number of recombinant sequences. The disproportionate geo-graphic distribution of recombinant DENV sequences likely reflects both the higher transmission intensity and the greater sampling effort in hyperendemic regions of Asia (Gubler, 2011; Bhatt et al., 2013; Messina et al., 2014). The detection of recombinants across four continents reinforces that recombination in DENV is global, and reinforces the need for surveillance in under-sampled regions.

Recombination events were distributed unevenly across the DENV genome: structural genes presented fewer recombination events (*C* : 0, pr: 2, *M* : 1, *E* : 8) than non-structural genes (*NS1* : 8, *NS2* : 12, *NS3* : 19, *NS4A*: 9, *NS4B*: 11, *NS5* : 17). The predominance of recombination in non-structural genes suggests that such *loci* may tolerate genetic exchange without compromising viral fitness.

Most recombination events were detected in *NS3, NS5*, and *NS2*, which are essential for viral replication (Sinha et al., 2024). The NS3 protein, together with its co-factor NS2B, mediates cleavage of the viral polyprotein and host proteins that could otherwise impair dengue infection (Lee et al., 2021). The NS5 protein plays a key role in RNA synthesis and mRNA capping and has been associated with suppression of the host antiviral response (Henderson et al., 2011; Tay et al., 2013). The high frequency of recombination in these genes is relevant as alterations in such genes could influence replication efficiency, viral fitness, and adaptability.

We identified eight recombination events within the *E* gene, which codes a protein involved in receptor binding. Recombination in *E* is clinically relevant because it mediates host cell entry and contains major neutralizing epitopes (Uno and Ross, 2018). Genetic exchange involving *E* has also been reported as relevant for live-attenuated vaccine strategies, as it may facilitate recombination between vaccine strains and circulating viruses, with potential effects on host specificity or virulence (Seligman and Gould, 2004).

The absence of recombination events in the capsid gene is noteworthy yet biologically consistent. The C protein is essential for nucleocapsid assembly and genome packaging, and this protein sequence is highly conserved because even minor disruptions can compromise viral structure and infectivity (Martín et al., 2002). These results align with previous studies reporting that recombination within the capsid gene is rare (Martín et al., 2002; Bouslama et al., 2007; Simmonds, 2006; Simmonds and Welch, 2006), reinforcing that strong functional constraints limit genetic exchange in regions critical for viral integrity.

### 3.3. Recombinant sequence analysis

All 51 recombination events with identified minor and major parents, as well as known breakpoints, were ana-lyzed for SNPs. Each recombinant sequence was compared to its minor parent in the recombinant region and to its major parent in the non-recombinant region. Recombinant sequences presented few SNPs relative to their minor parents in recombinant regions and few SNPs relative to their major parents in non-recombinant regions (Figure 1). In recombinant regions, SNP percentages ranged from 0% to 36.7% (mean 2.38%; SD 5.65%). In non-recombinant regions, they ranged from 0.04% to 30.29% (mean 2.19%; SD 4.27%). The low SNPs of recombinants compared to their parents are consistent regardless of gene, genotype, or serotype. The low SNP counts between recombinants and their parents correspond to a high sequence identity within each region, supporting the inference that such recombinant regions were inherited from those sequences.

**Figure 1.**
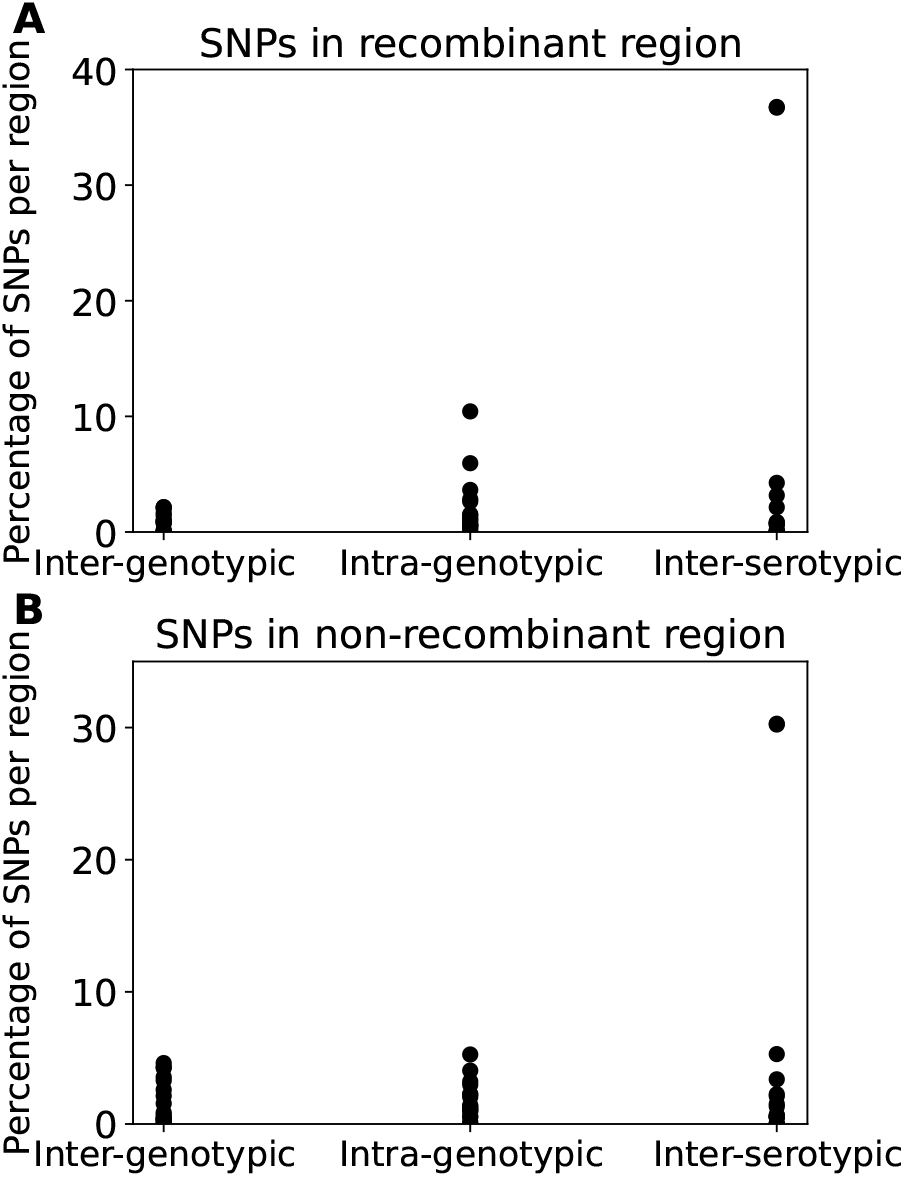
SNP divergence: recombinant vs. parental. (A) SNP percentages in recombinant regions relative to minor parents and (B) non-recombinant regions relative to major parents.

Seven recombination events were selected as case studies for detailed SNP and similarity analyses (Figures S2 to S8). In all cases, SNP distribution within recombinant regions was consistent with the pattern described above and was accompanied by higher pairwise similarity to the minor parent in these regions. For example, the intra-genotypic event involving recombinant JQ922559, minor parent JF262783, and major parent MW945600, all classified as DENV-4 I, spans the *E* gene between alignment positions 2461 and 3262 (Figure 2). Within the 756 nt recombinant region, the recombinant sequence differed from the minor parent by six SNPs (0.79%) and from the major parent by 63 SNPs (8.33%).

**Figure 2.**
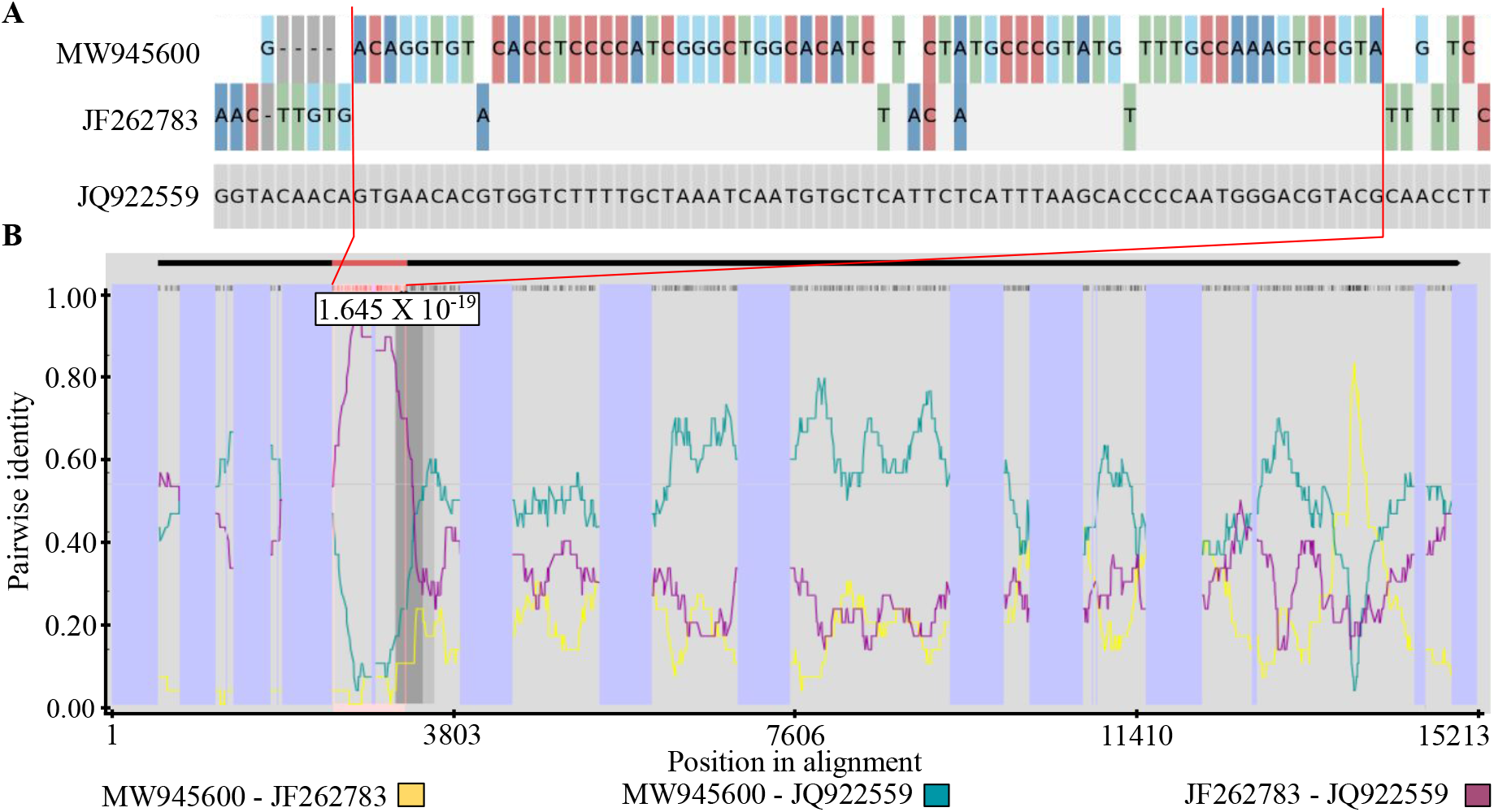
Sequence comparison among recombinant JQ922559, minor parent JF262783, and major parent MW945600 in the 2461–3262 region (red lines), spanning the *E* gene. (A) SNP analysis using SNIPIT. (B) RDP5 pairwise sequence similarity. Nucleotide colors: A (dark blue), C (red), T (green), G (light blue).

Interestingly, we depict in the case study 2 a recombination event with no minor parent identified (Figure S3). The recombinant presented a 26.45% SNP ratio in the recombinant region compared to the major parent. Given that sequences within the same DENV genotype diverge no more than 6% (de Souza et al., 2024), this high divergence could suggest that the minor parent belongs to a genetically distant lineage. A prior study has reported evidence that unidentified minor parents indicate recombination with unsampled, divergent lineages, as reported for MERS-CoV (Tolentino et al., 2024).

### 3.4. Literature search for the detected recombination events

The results of the literature search are summarized in Table S7. A total of 17 related studies were identified. Of the 53 recombinant sequences detected in this study, 27 had been previously analyzed for recombination. Furthermore, 26 of the 66 recombination events had already been reported (see Table S7 for references). Most studies (13) used RDP, typically as a standalone program (11) and less frequently in combination with GARD (Kosakovsky Pond et al., 2006) (2) or SimPlot (Lole et al., 1999) (1).

Most DENV genomes were classified as recombinant (17), while only three were classified as non-recombinant (Table S7). Notably, sequences GQ398257, FJ898399, JQ922553, JQ922559, and KY937189 received conflicting classifications across studies, being described as recombinant in some analyses and non-recombinant in others. Comparison of minor and major parent assignments revealed limited concordance between studies. Only one recombination event from sequence JQ922552 reported in related work matched both parent sequences identified in our analysis, whereas other events differed in one or both parent assignments (see Table S7 for references). Such discrepancies may reflect the dependence of recombination detection and parent inference on dataset composition and methodological choices (Jaya et al., 2023).

A total of 29 sequences are considered newly described recombinants in this work (Table 2; IDs in bold). These 29 sequences include 26 that were not examined in previous studies and three (JN638572, JN903578, and KF184975) that had been analyzed but not identified as recombinants. These 29 sequences were collected between 1993 and 2020 and are distributed across the four DENV serotypes: seven DENV-1, 16 DENV-2, three DENV-3, and three DENV-4.

### 3.5. Phylogenetic analysis of recombinant DENV genomes

The tree was rooted using Zika virus (MH882548) and West Nile virus sequences (HM147823, GQ903680, and JX041632) as outgroup taxa. The monophyly of DENV serotypes 1, 2, 3, and 4 was recovered (Figure 3). The DENV-1 and DENV-3 clades formed sister groups, as previously reported (Phadungsombat et al., 2018; Shrivastava et al., 2018).

**Figure 3.**
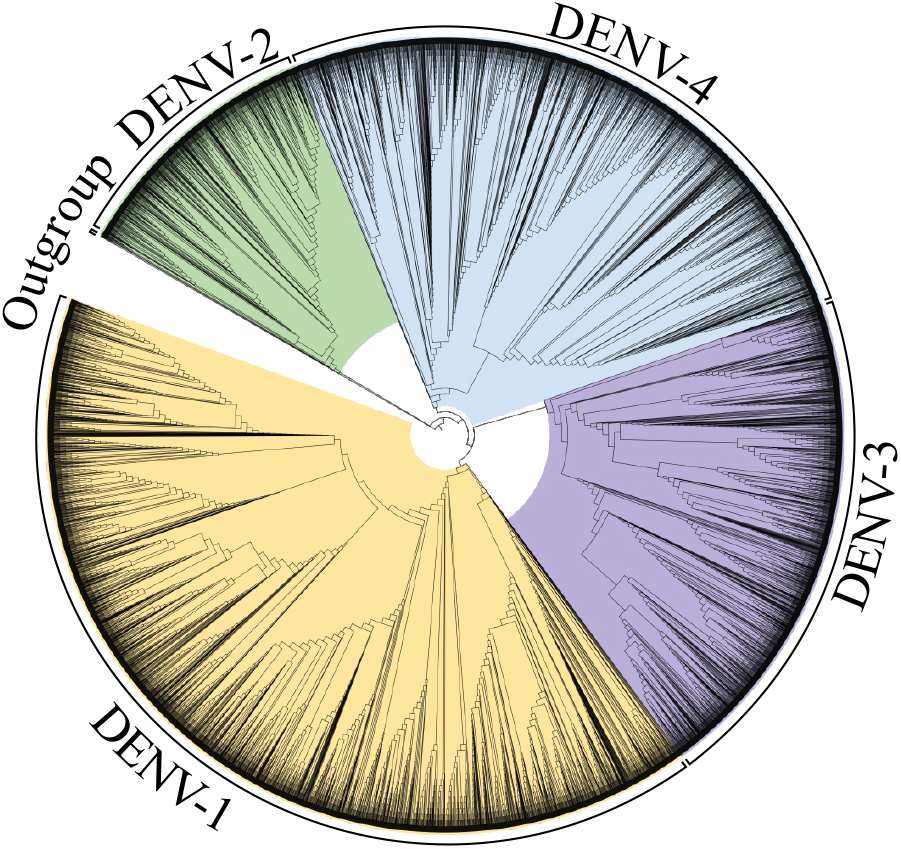
ML phylogenomic cladogram of DENV. Ingroup: 6,638 sequences; Outgroup: 1 ZIKV, 3 WNV. Colors: DENV-1 (yellow), -2 (green), -3 (purple), -4 (blue).

A weak positive correlation was observed between the proportion of recombinant terminals per clade and branch length (Spearman *ρ* = 0.087; *p* = 1.61 *×*10^−23^). The null hypothesis of equal branch length distributions between recombinant and non-recombinant terminals was not rejected at a significance threshold of 0.05 (Mann–Whitney U test, *p* = 4.35 *×* 10^−28^). Despite the statistically significant but weak correlation, recombinant sequences tended toward longer terminal branches, consistent with additional substitutions required to accommodate their phylogenetic placement (Schierup and Hein, 2000; Lukashev et al., 2003; Lemey et al., 2009; Didelot and Wilson, 2015). Given the weakness of the association, branch length alone cannot serve as a reliable indirect test for recombination.

Recombinant sequences were positioned closer to their major parents than to their minor parents in the phylogenetic tree. This difference was statistically significant when comparing the distributions of phylogenetic dis-tances between recombinants and their major parent clades versus their minor parent clades (*p* = 8.9*×* 10^−6^; test statistic = 155). The closer proximity of recombinants to their major parents likely reflects their larger genomic contribution, resulting in higher sequence identity. In contrast, distances between recombinants and their minor parents were more variable, consistent with their smaller genomic contribution and lower sequence identity in the whole-genome level.

Removing individual recombinant sequences for tree inference did not increase topological disruption, despite previous reports suggesting that recombinant sequences may affect tree topology (Posada et al., 2002; Lam et al., 2010). To assess this, trees generated after excluding recombinant terminals were compared with those excluding non-recombinant terminals. No significant difference was observed between these trees (two-tailed Mann–Whitney test, *p* = 0.897797). This result likely reflects the relatively short length of recombinant regions and the high sequence similarity among DENV genomes, which may limit the introduction of conflicting phylogenetic signals.

### 3.6. Performance evaluation of RECOSIM

All combinations of RECOSIM clustering strategies and parameter values were evaluated using a dataset of 160 simulated alignments with recombination. In total, 48 combinations were tested, comprising the three clustering strategies and the following parameter values: overlap_threshold (o) = 20 or 30, identity_threshold (i) = 0.90 or 0.95, min_samples (m) = 3 or 5, and epsilon (e) = 20 or 30. Performance metrics for each clustering strategy and parameters are summarized in Table S8. Overall, hit ratios ranged from 15.63% to 33.13%, while fail ratios ranged from 5.6% to 18.2%. The best-performing combination used the regularClusters.py implementation with o = 30, i = 0.95, e = 30, and m = 3, achieving a hit ratio of 33.13% and a fail ratio of 6.56%.

The performance of RECOSIM, using the optimal combination, and RDP5, using the previously described parameters, evaluated 160 simulated and 140 empirical alignments. Detection performance was measured by precision and sensitivity. RECOSIM achieved higher precision, with 93.44% in simulated data and 98.11% in empirical data, compared to 80% and 39.33% for RDP5 (Table S9). This indicates that recombination events detected by RECOSIM are more likely to be correct. Sensitivity values were lower and more similar between the tools. RECOSIM reached 35.62% and 37.14% in simulated and empirical data, respectively. RDP5 showed lower sensitivity in simulated data (22.5%) but higher sensitivity in empirical data (42.14%).

These trade-offs reflect distinct methodological priorities. RECOSIM emphasizes high-confidence detection with fewer false positives, whereas RDP5 detects more events in some scenarios at the cost of reduced specificity, consistent with previous reports (Martin et al., 2021). Both tools exhibited relatively low sensitivity below 45%, primarily due to high false-negative rates exceeding 60% in all cases. These results indicate that a substantial proportion of recombination events remains undetected, highlighting the difficulty of identifying subtle or complex recombination patterns.

## Acknowledgements

We gratefully acknowledge the institutional and computational support provided by the LNCC (Brazil) and the UNC Charlotte (USA), including the Center for Computational Intelligence to Predict Health and Environmental Risks (CIPHER), the Department of Bioinformatics and Genomics, the College of Computing and Informatics, and the University Research Computing group, which made this research possible. We also acknowledge CAPES and CNPq for financial support.

## Supplementary materials

Supplementary materials for this article are available via Zenodo at https://doi.org/10.5281/zenodo.20059036.

## Data availability

All data analyzed in this study and all supplementary material are available via Zenodo at https://doi.org/10.5281/zenodo.20059036. Further information is available from the authors upon request.

## CRediT authorship contribution statement

**HPO:** Software, Validation, Formal analysis, Investigation, Data Curation, Writing – original draft, Writing – review & editing, Visualization. **DJM:** Conceptualiza-tion, Methodology, Software, Validation, Formal analysis, Investigation, Resources, Data Curation, Writing – review & editing, Visualization, Supervision, Project administration, Funding acquisition. **PPO**: Formal analysis, Writing – review & editing. **KO:** Conceptualization, Methodology, Validation, Formal analysis, Investigation, Resources, Data Curation, Writing – original draft, Writing – review & editing, Visualization, Supervision, Project administration, Funding acquisition.

